# How a single sperm protein recognizes highly divergent egg coat components

**DOI:** 10.64898/2026.07.29.741504

**Authors:** Shunsuke Nishio, Han Wang, Luca Jovine

**Author notes:** S.N. and H.W. contributed equally to this work.

## Abstract

For successful fertilization, sperm must recognize and penetrate the egg coat, called vitelline envelope (VE) or, in mammals, zona pellucida (ZP). In abalone, sperm acrosomal protein lysin dissolves the VE by binding the N-terminal region of egg coat subunit VERL, encompassing 22 ZP-N domains (VR1-22), but how sperm engages other VE components remains unclear. Here we report free and lysin-bound structures of VEZP14-N1, the single ZP-N repeat of VERL paralog VEZP14, together with biochemical and biophysical data. Despite ∼20% sequence identity to VRs, VEZP14-N1 presents a structurally equivalent surface to lysin, indicating that conserved interaction geometry underlies recognition. Binding is asymmetric: lysin is conformationally rigid, whereas VEZP14-N1 requires homodimer dissociation and induced-fit rearrangements to bind — kinetic costs that limit affinity to an intermediate level, consistent with a decoy role sequestering lysin from VERL. These findings illuminate how one sperm protein binds divergent egg coat targets, with implications for ZP-N repeat function in vertebrate gamete recognition.

## Introduction

Fertilization requires sperm to interact with the egg coat extracellular matrix and penetrate it to ultimately fuse with the oocyte plasma membrane (1). In the marine gastropod abalone, VE contact triggers sperm acrosomal exocytosis, releasing lysin, which locally dissolves the egg coat in a species-specific way by binding VERL’s VR repeats through a non-enzymatic mechanism thought to reflect electrostatic repulsion between lysin-bound VERL subunits (2–5). Like vertebrate ZP components, which can also contain N-terminal ZP-N domains involved in egg coat architecture and gamete recognition (5–7), abalone VE subunits polymerize using a C-terminal ZP module (4, 8). However, despite its higher complexity, the abalone VE contains only one other subunit, VEZP14, which harbors an N-terminal ZP-N domain that also binds lysin (9). Via this interaction, VEZP14 was proposed to act as a molecular decoy reducing the effective concentration of free lysin able to bind VERL (10). Although we previously determined the structural basis of the interaction between lysin and different VRs (5), how lysin can also engage VEZP14, a highly sequence divergent paralog of VERL, remains undefined.

## Results and Discussion

Mature VEZP14 consists of six 19-residue T/S-rich repeats (encoded by exons 2–7) predicted to form a highly N/O-glycosylated solenoid; a short linker (exon 8); the N-terminal ZP-N domain (ZP-N1; exon 9); a stalk region consisting of T_3–7_P repeats also predicted to be heavily O-glycosylated; and a ZP module (exon 10) (Fig. 1A). A construct encompassing red abalone linker/ZP-N1 residues G146–G274 was well expressed in mammalian cells and sufficient for pulling down conspecific lysin, with an additional cysteine (C189) outside the canonical ZP-N disulfides (C_1_-C_4_, C_2_-C_3_) (5) being dispensable (Fig. 1B). For subsequent studies we thus used a C189Q mutant, henceforth referred to as VEZP14-N1. Size-exclusion chromatography with multi-angle light scattering (SEC-MALS) shows that purified VEZP14-N1 is a noncovalent homodimer, which dissociates upon incubation with lysin to form a 1:1 heterodimer (Fig. 1C).

**Fig. 1.**
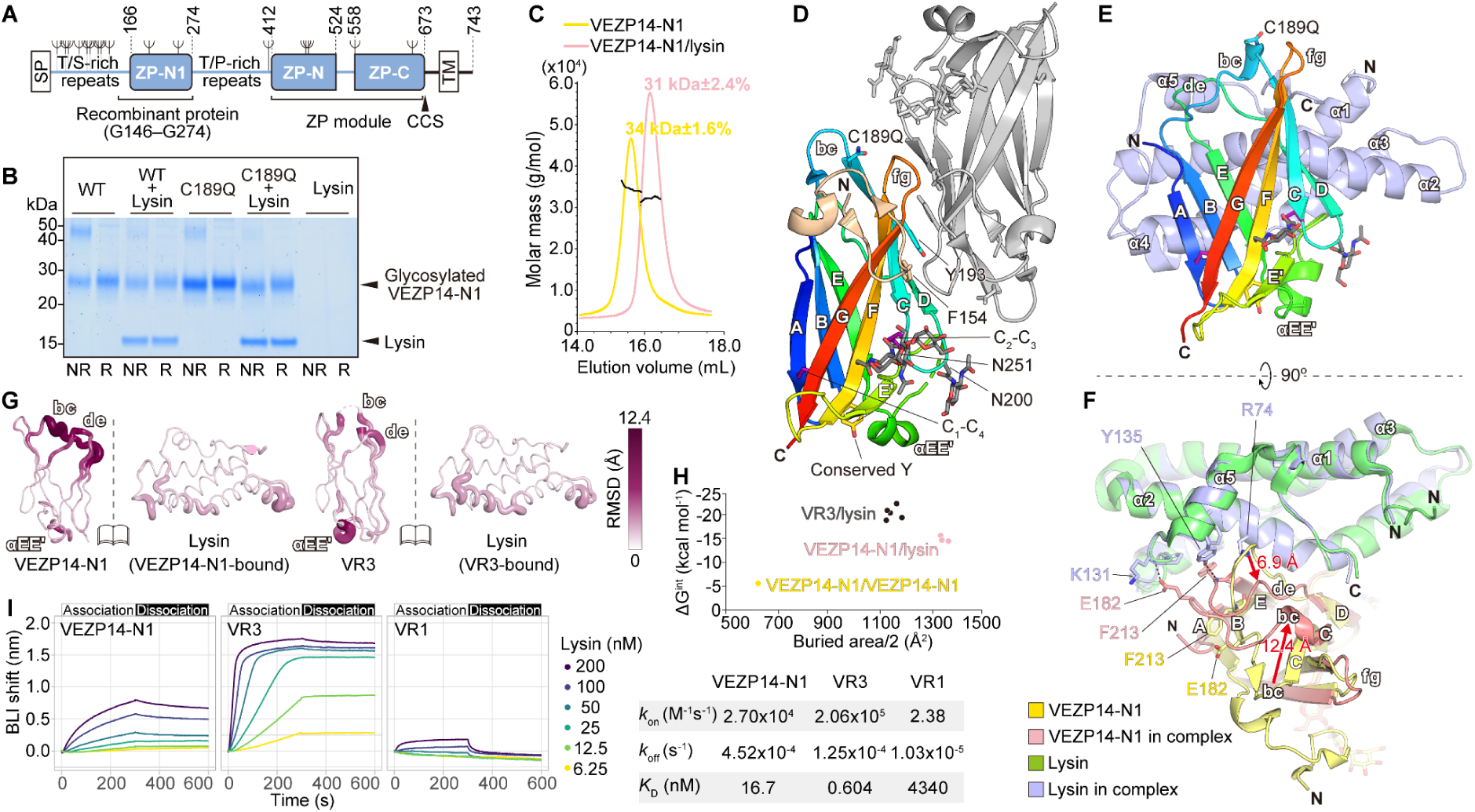
Crystal structures of VEZP14-N1 and its complex with lysin. (A) The secreted ectodomain of VEZP14 (blue) is produced from a type-I transmembrane precursor upon signal peptide (SP) cleavage and processing at a consensus cleavage site (CCS) preceding the C-terminal transmembrane domain (TM); ψ, predicted N-glycosylation sites. (B) Pull-down of untagged lysin co-transfected with His-tagged WT VEZP14 G146–G274 or C189Q mutant (VEZP14-N1), analyzed under non-reducing (NR) or reducing (R) conditions. (C) SEC-MALS of purified deglycosylated VEZP14-N1 and VEZP14-N1/lysin. (D) Structure of free, homodimeric VEZP14-N1. The ZP-N1 domain of one subunit is colored with a rainbow gradient from blue (N-terminus) to red (C-terminus), with the short N-terminal linker colored wheat. The second subunit is gray. (E) Structure of VEZP14-N1/lysin, with VEZP14-N1 colored as in panel D and lysin in light blue. (F) Structural comparison of VEZP14-N1, lysin (PDB ID: 5II8) and their complex. Red arrows highlight the rearrangements of VEZP14-N1 bc and de loops upon lysin binding. (G) VEZP14-N1 and VR3 residue RMSDs between unbound and lysin-bound states. (H) Solvation free energy gains upon interface formation (ΔG^int^) and surface buried areas. (I) BLI sensorgrams of lysin and VEZP14-N1, VR3, or VR1.

Although AlphaFold predictions of VEZP14-N1 have very low confidence (mean predicted Local Distance Difference Test (pLDDT): 30.9 (11)), we could use a model ensemble to solve the 1.54 Å resolution crystal structure of the free protein by molecular replacement followed by simulated annealing refinement. The resulting experimental model was then used to determine the structure of the VEZP14-N1/lysin complex, in two different crystal forms diffracting to 1.5 and 1.84 Å resolution, respectively (Fig. 1D, E).

VEZP14-N1 adopts an immunoglobulin-like ZP-N fold, with β-strand D (βD) belonging to β-sheet 2 as observed in VRs (5). Conserved D156, T192, Y193, Q195, T202, N259 and R262 in the fg loop-containing half of this sheet mediate VEZP14-N1 homodimerization, which is also stabilized by N-terminal linker F154 (Fig. 1D). Although the resulting dimer is different from that of VR2 (5), both VEZP14 and VERL are recognized by lysin through βA, βB, βD and βE (Fig. 1E, 2A). Comparison of the free and bound structures explains why lysin dissociates the VEZP14-N1 homodimer: two lysin molecules could not bind simultaneously without sterically clashing. The comparison also reveals that, whereas the lysin/VR interaction is essentially rigid-body (average free/bound RMSDs of 0.7 Å over 124 Cα atoms for lysin and 1.6 Å over 102 Cα for VRs (5)), binding of lysin to VEZP14-N1 involves substantial induced fit in the latter (3.4 Å over 105 Cα). In particular, VEZP14-N1 loop bc shifts by 12 Å, positioning the side chain of E182 to hydrogen bond to the backbone amide of lysin K131, whereas loop de moves by 7 Å, enabling main-chain atoms of F213 to hydrogen bond with the side chains of lysin R74 (α-helix 2) and Y135 (α-helix 5) (Fig. 1F, G).

**Fig. 2.**
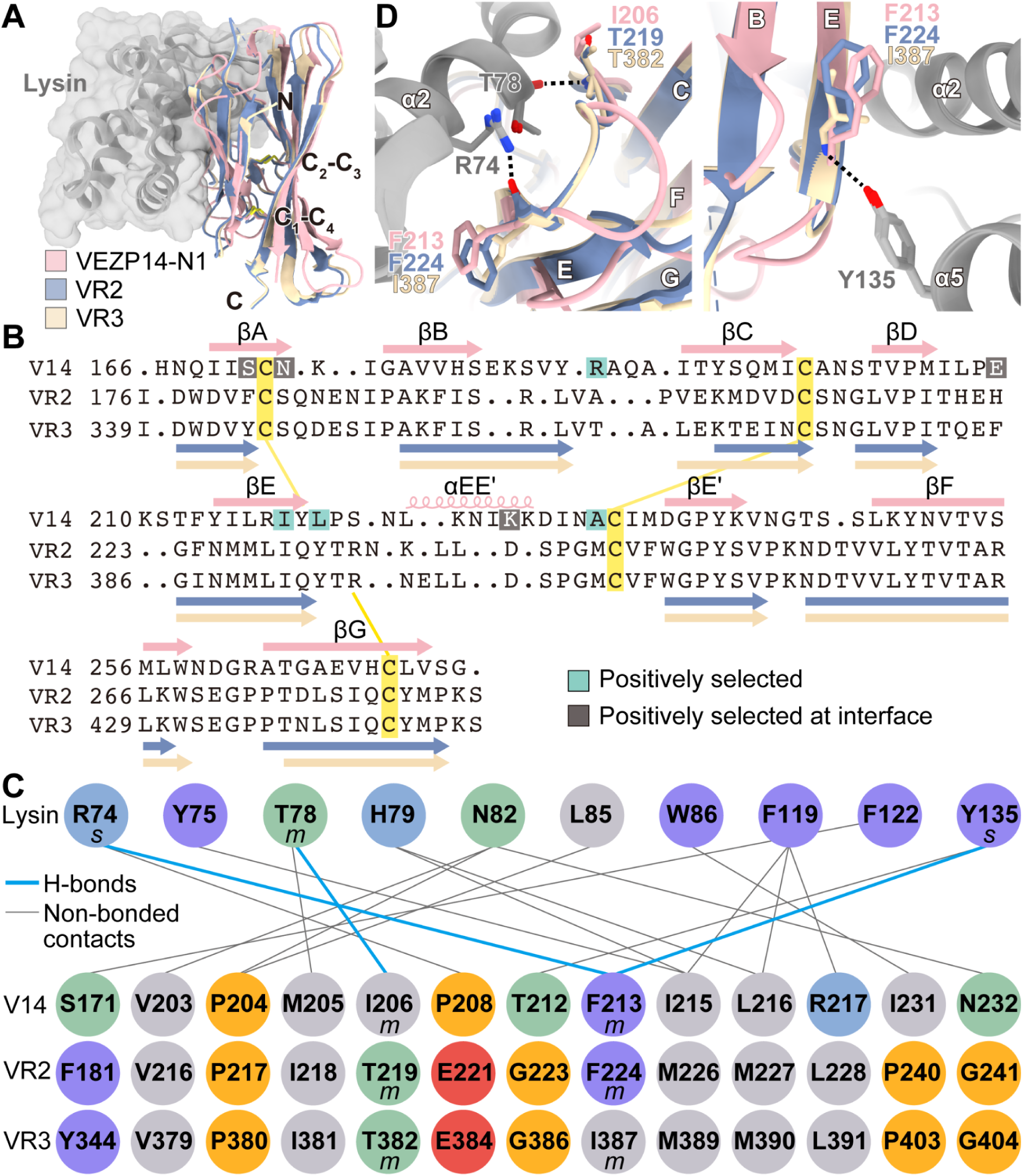
Structural comparison of VEZP14-N1/lysin with VR/lysin complexes. (A) Structural superposition of VEZP14-N1/lysin, VR2/lysin (PDB ID: 5MR3) and VR3/lysin (PDB ID: 5IIA). Here and in panel D, lysin is gray. (B) Structure-based sequence alignment of VEZP14-N1 (V14) and VRs. (C) Interactions found in all copies of VEZP14-N1/lysin, VR2/lysin and VR3/lysin. s, side chain; m, main chain. (D) Hydrogen bonds conserved in all complexes.

Computational analysis indicates that VEZP14-N1/lysin is more stable than the VEZP14-N1 homodimer but less stable than the complex between lysin and a high-affinity repeat representative of VR3-22 (VR3; Fig. 1H). Accordingly, measurements of binding kinetics by biolayer interferometry (BLI), using VR1 and lysin (whose interaction cannot be detected in solution (5)) as a negative control, show that VEZP14-N1/lysin has a 30-fold higher dissociation constant (*K*_D_) than VR3/lysin, while exhibiting a lower association rate (*k*_on_) but similar dissociation rate (*k*_off_) (Fig. 1I). This is consistent with lysin binding depending on both VEZP14-N1 homodimer dissociation and induced-fit rearrangements, whose combined entropic cost accounts for the lower *k*_on_ and weaker affinity of VEZP14-N1/lysin relative to VR3/lysin (Fig. 1G), despite a larger buried interface in the former (Fig. 1H) (12).

Although VEZP14-N1 is only 19.8% and 18.9% sequence identical to VR2 and VR3, respectively, structural alignment of the three complexes with lysin (Fig. 2A, 2B) shows that they share 18 interactions (Fig. 2C), with lysin R74, T78 and Y135 forming hydrogen bonds to structurally equivalent main chain atoms of each egg coat counterpart (Fig. 2D). Thus, lysin recognizes conserved geometric and surface features rather than sequence, providing a molecular explanation for how a single sperm protein can bind multiple egg coat targets of highly divergent sequence.

VEZP14-N1 and VRs are paralogous ZP-N domains that, despite very low sequence identity, both engage the same rigid lysin surface — VRs through rigid-body docking, VEZP14-N1 through homodimer dissociation and induced fit. The resulting affinity of VEZP14, intermediate between non-binding VR1 and high-affinity VR3, supports a decoy role (10) because — considering the relative abundance of VERL and VEZP14 (9) — binding as tightly as VR3 would allow VEZP14 to outcompete VERL itself.

Interestingly, Lys to Pro substitution of positively selected lysin residue 131 in green abalone would preclude the aforementioned hydrogen bond to VEZP14 E182; on the other hand, lysin’s hypervariable, positively selected N-terminus remains disordered in the VEZP14-N1 complex, as it does with VRs (5), indicating that this region is not co-opted for binding even by a paralog that engages lysin through a markedly different mechanism. From the VEZP14 side, four out of eight positively selected residues (10) map to the lysin interface (Fig. 2B), confirming that selection specifically targets this surface; however, modeling of corresponding substitutions found in other *Haliotis* species suggest that they would not negatively affect (and in some cases may even enhance) binding to red abalone lysin, arguing against selection for increased conspecific affinity at this interface and leaving open what other property these residues may be under selection for. Together, these results shed light on the complexity of gamete recognition and provide a structural basis for the decoy function of VEZP14 first proposed from evolutionary analyses (10), while underscoring that the forces shaping its evolution remain incompletely understood.

## Materials and Methods

Proteins were expressed in HEK293 cells. Diffraction data were collected at beamlines I04, Diamond Light Source (United Kingdom), and ID30A-3, European Synchrotron Radiation Facility (France). Binding kinetics were determined by BLI. A full description of the methodology is detailed in SI Appendix, Materials and Methods.

## Data, Materials, and Software Availability

Structure factors and model coordinates are deposited in Protein Data Bank (PDB) [31BB (13), 31BC (14), 31BD (15)]. All study data are included in the article and/or *SI Appendix*.

## Acknowledgments

This research was supported by Swedish Research Council grants 2020-04936 and 2024-05336, and Knut and Alice Wallenberg Foundation grant 2018.0042 (L.J.). We thank Hamed Sadat, Dirk Fahrenkamp and Benjamin Wiseman for early involvement in the project; Masaya Hane and Chihiro Sato for help with BLI; Javier Delgado for advice on FoldX; Alena Stsiapanava and Daniele de Sanctis for discussions.

## Author contributions

S.N. and L.J. designed research; S.N. and L.J. performed research; S.N., H.W., and L.J. analyzed data; and H.W., S.N., and L.J. wrote the paper.

## Supplementary Materials and Methods

### DNA constructs

A gene encoding the ZP-N1 domain of *Haliotis rufescens* VEZP14 (corresponding to residues G146–G274 of Uniprot entry D0EL52, with a C189Q substitution to pre-empt spurious intermolecular disulfide formation) was synthesized (ATUM) and cloned into mammalian expression vector pLJ6 (1), in frame with sequences encoding an N-terminal chicken Crypα signal peptide and a C-terminal 8His-tag. An equivalent expression vector for wild-type (C189) VEZP14-N1 was then generated by Seamless Ligation Cloning Extract (SLiCE)-mediated mutagenesis (2). VRs and lysin were produced using mammalian expression vectors pHLsec3-VR3+, pHLsec3-VR1+ and bacterial expression vector pJexpress411-LYSIN-02, respectively, as previously documented (3).

### Protein expression and purification

For biochemical experiments, recombinant VEZP14-N1 and VRs were expressed in HEK293T cells; for SEC-MALS analysis and crystallization, VEZP14-N1 was expressed in HEK293S cells. 25 kDa branched PEI (Sigma-Aldrich) was used for transient transfection, following an established protocol (4). Four days after transfection, the conditioned medium was collected by centrifugation at 6000 x *g* for 15 min at 4°C and filtered through 0.8 µm and 0.2 µm filters. The supernatant was adjusted to IMAC binding buffer conditions (20 mM Na-HEPES pH 7.8, 150 mM NaCl, 10 mM imidazole) and Ni Sepharose excel (GE Healthcare) was then added at a ratio of 1:100 (v/v). After overnight incubation at 4°C, beads were packed into a column, washed with 100 volumes of IMAC binding buffer, and proteins were eluted with 20 mM Na-HEPES pH 7.8, 150 mM NaCl, 500 mM imidazole. In the case of the VEZP14-N1 expressed in HEK293S, high-mannose glycans were trimmed with Endoglycosidase H (Endo H; 1:10–1:20 mass ratio) for 3 h at room temperature before imidazole elution. IMAC eluates were concentrated using 10 kDa cutoff centrifugal filters (Amicon) and further purified by SEC at 4°C using an ÄKTA FPLC system (GE Healthcare) and a Superdex 75 Increase 10/300 GL column (GE Healthcare), pre-equilibrated with 20 mM Na-HEPES pH 7.8, 150 mM NaCl. Peak fractions were pooled, concentrated and stored at −80°C.

After expression in *E. coli* BL21 Star (DE3) (Thermo Fisher Scientific) co-transformed with pRARE (Novagen), lysin was refolded from inclusion bodies and purified as described previously (3). Peak fractions from ion-exchange chromatography using a 5 mL HiTrap CM Sepharose FF column (GE Healthcare) were collected, dialyzed against 50 mM sodium acetate pH 5.5, concentrated to 20 mg/mL and stored at −80°C.

To form the VEZP14-N1/lysin complex, proteins were incubated at 1:1.5 ratio (w/w) overnight at 4°C. After clearing the sample by centrifugation at 17,000 x g for 40 min at 4°C, the complex was purified by SEC, using the same column and conditions described above. Peak fractions were pooled and concentrated to 30 mg/mL.

### Protein crystallization

Crystallization trials were performed at 20°C using the sitting-drop vapor-diffusion method, either with a mosquito crystallization robot (TTP Labtech) or manually. Diamond-shaped crystals of VEZP14-N1 were obtained by mixing 1 µL protein solution (2.7 mg/mL) with 1 µL reservoir solution containing 0.1 M Tris-HCl pH 8.5, 25% (w/v) PEG 4000.

For the VEZP14-N1/lysin complex, two crystal forms were obtained. The P1 crystal form was obtained by mixing 0.1 µL protein solution (23–28 mg/mL) with 0.1 µL reservoir solution containing 0.2 M (NH_4_)_2_SO_4_, 0.1 M sodium acetate pH 4.6, 30% (v/v) PEG 2000 MME. The C2 crystal form was obtained by mixing 0.1 µL protein solution with 0.1 µL reservoir solution containing 1 M LiCl, 0.1 M citric acid pH 4.0, 20% (w/v) PEG 6000. Both forms appeared as clusters of thin plate-like crystals, which were separated using MicroLoops (MiTeGen) during harvesting and flash frozen in liquid N_2_ for data collection.

### X-ray diffraction data collection

All datasets were collected from single crystals at 100 K.

Trigonal datasets for free VEZP14-N1 were collected at a wavelength of 0.9795 Å at Diamond Light Source beamline I04 (5), using an EIGER2 X 16M detector (DECTRIS).

Triclinic and monoclinic data for the VEZP14-N1/lysin complex were collected at a wavelength of 0.9677 Å at beamline ID30A-3/MASSIF-3 of the European Synchrotron Radiation Facility (ESRF) (6), using an EIGER X 4M detector (DECTRIS).

### Data processing and structure determination

All final datasets were processed using XDS (7).

For free VEZP14-N1, data collection statistics generated using phenix.table_one (8, 9) were: completeness 99.6 (99.3) %, multiplicity 6.5 (6.6), mean I/sigma(I) 12.9 (0.5), R_merge_ 0.058 (2.840), R_meas_ 0.064 (3.081), R_pim_ 0.025 (1.178), CC_1/2_ 1.00 (0.57), CC* 1.00 (0.85) (with the values in parentheses being for the highest resolution shell, 1.57–1.54 Å).

For the *P*1 crystal form of VEZP14-N1/lysin, statistics were: completeness 96.5 (93.0) %, multiplicity 6.1 (5.3), mean I/sigma(I) 5.5 (0.8), R_merge_ 0.165 (1.534), R_meas_ 0.180 (1.700), R_pim_ 0.071 (0.725), CC_1/2_ 0.99 (0.37), CC* 1.00 (0.74) (with the values in parentheses being for the highest resolution shell, 1.52–1.50 Å).

For the *C*2 crystal form of VEZP14-N1/lysin, statistics were: completeness 98.3 (98.8) %, multiplicity 3.6 (3.7), mean I/sigma(I) 4.7 (0.8), R_merge_ 0.170 (1.550), R_meas_ 0.201 (1.813), R_pim_ 0.105 (0.933), CC_1/2_ 0.99 (0.40), CC* 1.00 (0.76) (with the values in parentheses being for the highest resolution shell, 1.89–1.84 Å).

The structure of free VEZP14-N1 was solved in space group *P*3_2_21 by molecular replacement (MR) with Phaser (10), using as search model an ensemble of 5 predictions of residues A147–G274[C189Q] generated by a local installation of AlphaFold 2 (11) (mean pLDDT per model: 42.3–79.2). For MR, AlphaFold-generated pLDDT values were converted to pseudo-B-factors by subtracting each residue’s pLDDT score from 100, and Phaser was run with an expected root-mean-square (RMS) coordinate error of 1.1 Å. This produced a single solution with one molecule per asymmetric unit (LLG 89, TFZ 11.9) that, however, resisted conventional refinement in phenix.refine (12) (R_free_ 0.54). Inclusion of simulated annealing and a larger number of macro cycles (15) allowed refinement to converge to R_free_ 0.47, yielding an improved model that produced much more convincing solution statistics when used as input for a second run of Phaser (LLG 2503, TFZ 34.0) and subsequent autobuilding with PHENIX AutoBuild (13) (R_work_ 0.28, R_free_ 0.30). After several cycles of manual rebuilding in Coot (14) alternated with refinement, geometry and all-atom contacts of the protein model (R_work_ 0.20, R_free_ 0.23) were validated using MolProbity (15): Ramachandran favored/allowed/outliers: 97.5/2.5/0.0 %; Rama distribution Z-score: 0.60 ± 0.79; favored/poor rotamers: 98.2/0.0 %; MolProbity score: 0.98; clashscore 1.47. Structure validation of carbohydrates attached to VEZP14 N200 and N251 in this and the other structures was carried out using Privateer (16). Consistent with the difficulty of the initial refinement, the final experimental model was significantly different from the AlphaFold predictions included in the MR search ensemble (RMS deviation 2.6–4.4 Å).

For determining the triclinic structure of the VEZP14-N1/lysin complex by MR, we used an initial dataset automatically processed by xia2/DIALS (17, 18) and two search models, a partially refined model of free VEZP14-N1 that included residues R162–L207[C189Q], T212–N223, C234–K241, L248–V272 and an ensemble of red abalone lysin structures (residues L29–M152) (3). Phaser produced a single solution, consisting of two copies of VEZP14-N1/lysin per asymmetric unit (LLG 3570, TFZ 18.8), which was rebuilt and refined to R_work_ 0.23, R_free_ 0.25 using PHENIX AutoBuild. Manual rebuilding in Coot and refinement with phenix.refine against the same dataset reprocessed with XDS (see previous paragraph) yielded a final model with R_work_ 0.17, R_free_ 0.20, which was validated as above (Ramachandran favored/allowed/outliers: 98.9/1.1/0.0 %; Rama distribution Z-score: 0.58 ± 0.34; favored/poor rotamers: 100.0/0.0 %; MolProbity score: 1.16; clashscore 3.66).

Separate ensembles of the refined VEZP14-N1 and lysin coordinates from the *P*1 structure were used to phase the *C*2 dataset of the same complex, resulting in a single high-scoring MR solution (LLG 5528, TFZ 67.1) with one copy of VEZP14-N1/lysin per asymmetric unit, which was rebuilt and refined to R_work_ 0.21, R_free_ 0.25 (Ramachandran favored/allowed/outliers: 99.6/0.4/0.0 %; Rama distribution Z-score: 1.31 ± 0.50; favored/poor rotamers: 98.5/0.0 %; MolProbity score: 0.85; clashscore 1.30).

### Small-scale pull-down experiments

Pull-down assays were performed following an established protocol (3), with expression plasmids mixed at a 1:1 DNA ratio for co-transfection of VEZP14-N1 and lysin in mammalian cells. Proteins recovered by pull-down were separated by SDS-PAGE and visualized using SimplyBlue SafeStain (Thermo Fisher Scientific).

### Size-exclusion chromatography-multi angle light scattering (SEC-MALS)

VEZP14-N1 or the VEZP14-N1/lysin complex (100–150 µg) were analyzed using an Ettan LC high-performance liquid chromatography (HPLC) system equipped with a UV-900 detector (Amersham Pharmacia Biotech; λ = 280 nm). This system was coupled to a miniDawn TREOS MALS detector (Wyatt Technology; λ = 658 nm) and an Optilab T-rEX dRI detector (Wyatt Technology; λ = 660 nm). Separation was performed at 20°C using a Superdex 200 Increase 10/300 GL column (GE Healthcare) at a flow rate of 0.5 mL/min. The mobile phase consisted of a seawater-like solution containing 468 mM NaCl, 10 mM KCl, 10 mM CaCl_2_, 28 mM MgSO_4_, 25 mM MgCl_2_, 20 mM Na-HEPES pH 7.8. Data was processed, and the weight-averaged molecular mass was calculated using ASTRA software (Wyatt Technology).

### Bio-layer interferometry (BLI) analysis

Lysin binding experiments were performed using an Octet K2 system (Sartorius) at the Institute for Glyco-core Research (iGCORE) of Nagoya University. VEZP14-N1, VRs were prepared at 0.01 mg/mL in a seawater solution (Jamarin Laboratory) and immobilized on Ni-NTA biosensors (Sartorius) overnight at 4°C. Association and dissociation measurements were performed in the same buffer. After baseline equilibration, immobilized sensor tips were immersed in lysin at concentrations ranging from 0 to 200 nM for 300 s, followed by dissociation in the buffer for another 300 s. Data were analyzed using a 1:1 binding model to determine the dissociation rate constant (*k*_off_), the association rate constant (*k*_on_), and the equilibrium dissociation constant (*K*_D_).

### Sequence-structure analysis

Sequence alignments were generated using ClustalO (19) and glycosylation sites were predicted using NetNGlyc (20) or NetOGlyc (21).

The fold of the N-terminal T/S-rich repeat region of VEZP14 was predicted using AlphaFold 2 and can also be observed in the AlphaFold Protein Structure Database prediction for full-length VEZP14 (22).

Structural models were visualized and analyzed using PyMOL (Schrödinger) and UCSF Chimera (23)/ChimeraX (24), which were also used to generate structural alignments (25).

Interfaces were analyzed using PISA (26), which was also used to calculate the ΔG^int^ and surface buried area of the VEZP14-N1 homodimer (PDB ID: 31BB), VEZP14-N1/lysin [two copies from the P1 crystal (PDB ID: 31BC), and one from the C2 crystal (PDB ID: 31BD)] and VR3/lysin (four copies from PDB ID: 5IIA, and one from PDB ID: 5IIB).

The effect of substitutions at red abalone VEZP14-N1 positively selected residues at the interface with lysin (S171, N173, E209 and K228 (27)) was evaluated with FoldX 5.1 (28). The BuildModel command of the program was used to introduce in combination the corresponding residues found in other *Haliotis* species for which a VEZP14 sequence is available (*H. corrugata*; *H. discus hannai*; *H. fulgens*; *H. sorenseni*; *H. kamtschatkana*; *H. walallensis* and *H. cracherodii*, with the *H. fulgens* and *H. kamtschatkana* complexes additionally modeled with the corresponding lysin substitution Y118F) into the experimental models of the *H. rufescens* VEZP14-N1/lysin complex; ΔΔG was calculated relative to the corresponding unmodified complexes. Calculations were performed in seawater-like conditions (--pH=8.1 --ionStrength=0.6 --temperature=288) on two independent complexes from the P1 crystal form (PDB ID: 31BC, chains A+B and C+D) and on the single complex from the C2 crystal form (PDB ID: 31BD), with prior structure repair (--command=RepairPDB) and taking into account experimental water molecules (--water=-CRYSTAL). This yielded 15 independent ΔΔG estimates per mutant (two quintuple replicates from P1, one quintuple replicate from C2) that were averaged. All VEZP14 species variant substitutions were predicted to have negligible effects on complex stability (ΔΔG −1.9 to −0.5 kcal/mol). As positive controls, the same protocol was applied to previously characterized mutations at the VR3/lysin interface (3), which were correctly predicted as strongly destabilizing (ΔΔG +2.7 to +9.3 kcal/mol).

